# In vivo CRISPR screens identify key modifiers of CAR T cell function in myeloma

**DOI:** 10.1101/2024.11.19.624352

**Authors:** Felix Korell, Nelson H. Knudsen, Tamina Kienka, Giulia Escobar, Celeste Nobrega, Seth Anderson, Andrew Y. Cheng, Maria Zschummel, Amanda Bouffard, Michael C. Kann, Sadie Goncalves, Hans W. Pope, Mitra Pezeshki, Alexander Rojas, Juliette S. M. T. Suermondt, Merle Phillips, Trisha R. Berger, Sangwoo Park, Diego Salas-Benito, Elijah P. Darnell, Filippo Birocchi, Mark B. Leick, Rebecca C. Larson, John G. Doench, Debattama Sen, Kathleen B. Yates, Robert T. Manguso, Marcela V. Maus

## Abstract

Chimeric antigen receptor (CAR) T cells are highly effective in hematologic malignancies. However, loss of CAR T cells can contribute to relapse in a significant number of patients. These limitations could potentially be overcome by targeted gene editing to increase CAR T cell persistence. Here, we performed *in vivo* loss-of-function CRISPR screens in BCMA-targeting CAR T cells to investigate genes that influence CAR T cell persistence, function and efficacy in a human multiple myeloma model. We tracked the expansion and persistence of CRISPR-library edited T cells *in vitro* and then at early and late timepoints *in vivo* to track the performance of gene modified CAR T cells from manufacturing to survival in tumors. The screens revealed several context-specific regulators of CAR T cell expansion and persistence. Ablation of *RASA2* and *SOCS1* enhanced T cell expansion *in vitro*, while loss of *PTPN2*, *ZC3H12A*, and *RC3H1* conferred early selective growth advantages to CAR T cells *in vivo*. Strikingly, we identified cyclin-dependent kinase inhibitor 1B (*CDKN1B*), a cell cycle regulator, as the most important factor limiting CAR T cell fitness at late timepoints *in vivo*. *CDKN1B* ablation increased BCMA CAR T cell proliferation and effector function in response to antigen, significantly enhancing tumor clearance and overall survival. Thus, our findings reveal differing effects of gene-perturbation on CAR T cells over time and in different selective environments, highlight *CDKN1B* as a promising target to generate highly effective CAR T cells for multiple myeloma, and underscore the importance of *in vivo* screening as a tool for identifying genes to enhance CAR T cell function and efficacy.

## Main

Chimeric antigen receptor (CAR) T cells have changed the landscape of treatment in hematologic malignancies, including B-cell leukemia, lymphoma, and multiple myeloma^1,2^. However, the currently approved CAR T cells are not curative for patients with relapsed or refractory multiple myeloma, and most patients eventually progress with disease that maintains expression of the target antigen. Patients often show a progressive loss of circulating CAR T cells, underscoring the need for additional research and modifications to enhance long-term CAR T efficacy and persistence^3–5^.

There are several genes and signaling pathways that could influence the persistence and efficacy of CAR T cells. One strategy to identify genetic modifications that confer increased persistence of CAR T cells is to use pooled loss-of-function genetic screens with CRISPR-Cas9-mediated genome editing. To date, genetic screens in T cells have largely been performed *in vitro* ^6–12^, where the selective pressure that is applied to the pool of gene-modified cells consists of either single or repetitive stimulation with antigen and the identification of cells that continue to produce cytokines or proliferate. However, *in vivo* models likely impose different selective pressures for persistence and fitness of CAR T cells, and might better reflect the physiological conditions that occur in patients.

Here, we develop and apply an *in vivo* pooled loss-of-function genetic screen to identify gene perturbations that can improve the expansion and therapeutic efficacy of BCMA CAR T cells against multiple myeloma as well as enhance their functionality and increase persistence.

## Longitudinal CRISPR screens in BCMA CAR T cells reveal genes that influence expansion and persistence *in vitro* and *in vivo*

We developed a CRISPR-based CAR T cell screening platform to discover genes that modify the expansion and persistence of human BCMA CAR T cells targeting MM.1S, a mouse xenograft model of human myeloma. To enable full recovery of a screening library *in vivo*, we designed a CRISPR single guide (sg)RNA library (the ‘Mario’ library) targeting 135 genes with known or proposed functions in T cells, with 8 sgRNAs targeting each gene to maximize statistical confidence in each gene hit (**Fig. 1a**). We included 100 intergenic sgRNAs as controls for a total library size of 1,080 sgRNAs (**Extended Data Table 1**). The Mario sgRNA delivery vector included a truncated LNGFR reporter (NGFR) and a dual sgRNA cassette in which the sgRNA library was cloned at position 1, downstream of the human U6 promoter, along with a TCRα constant (*TRAC*)-targeting sgRNA downstream of the human H1 promoter on the reverse strand in position 2 (**Extended Data Fig. 1a**).

**Figure 1.**
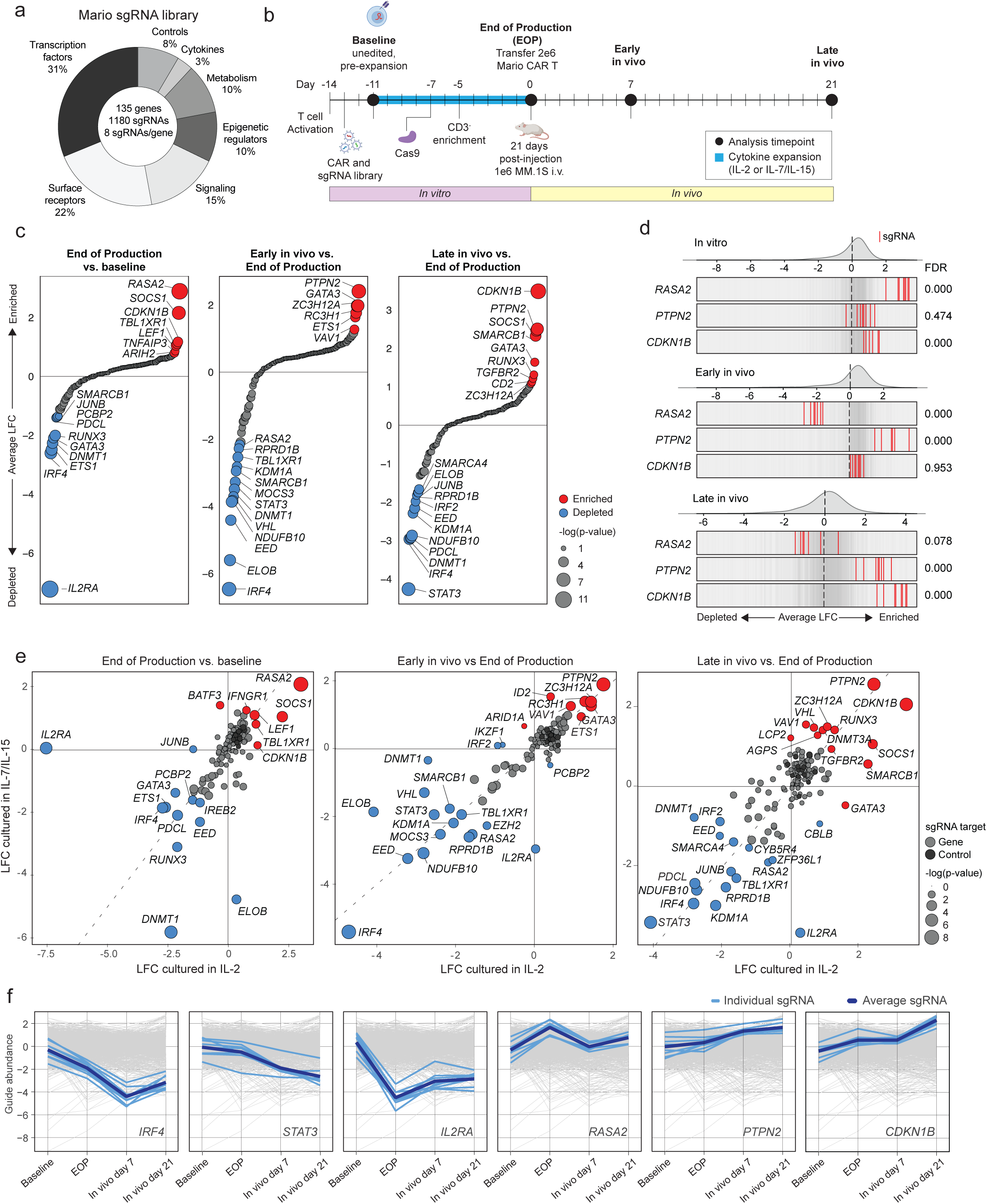
*In vivo* loss-of-function CRISPR screen identifies key regulators of CAR T cell function. **a.** Composition of genes targeted in the Mario library based on gene function. **b.** Diagram of Mario-CAR T cell production and screen workflow. T cells were activated with anti-CD3/CD28 microbeads, transduced one day later sequentially with the BCMA CAR and sgRNA library lentiviruses. A pre-electroporation sample was frozen down for analysis on day –11 (Baseline), and the remaining cells underwent Cas9 mRNA electroporation (day –7) and CD3 negative selection (day –5). Transduction efficiencies were assessed on day –4 and 0 at the end of production prior to cryopreservation. Mario-CAR T cells (2E6 double positive cells) were transferred into NSG mice bearing MM.1S multiple myeloma. At day 7 (early in vivo) or day 21 (late in vivo), mice were euthanized and a total marrow harvest was conducted (collecting femur, tibia, and spine) for isolation of Mario-CAR T cells. Mario-CAR T cells were produced from n=3 healthy donor T cells in IL-2 (ND216, ND99, ND106) and n=2 of the same donors for IL7/15 (ND106, ND216). **c.** Genes ranked by Log2(fold change) during in vitro manufacturing (end-of-production vs. baseline, left panel) and early in vivo (day 7 vs. end-of-production, middle panel) or late in vivo (day 21 vs. end-of-production, right panel). Enriched genes are shown in red and depleted genes are shown in blue with circle size corresponding to –log10 (FDR). n = 3 healthy donor T cells (ND216, ND99, ND106). **d.** Frequency histograms of enrichment or depletion of sgRNAs for *RASA2*, *PTPN2*, and *CDKN1B*, grouped by respective period. n = 3 healthy donor T cells (ND216, ND99, ND106). **e.** LFC comparison for gene knockout scoring between Mario-CAR T cells produced in IL-2 vs. IL-7/IL-15 at individual time points. n = 3 healthy donor T cells for IL2 (ND216, ND99, ND106) and 2 of the 3 donors for IL7/15 (ND106, ND216). **f.** Abundance of sgRNAs targeting individual genes across the entire screen workflow for Mario-CAR T cells produced in IL-2. **b** was created with Biorender.com. ND: normal donor. LFC: log fold change

Deletion of *TRAC* enabled the enrichment of successfully genome-edited cells through magnetic bead-based depletion of CD3^+^ cells (see **Methods**). The modular approach of a separate Mario sgRNA delivery vector enables the flexibility to evaluate CAR-or target-specific dependencies using a second lentiviral vector encoding the CAR (**Extended Data Fig. 1a**). Here, the BCMA CAR vector was synthesized based on idecabtagene vicleucel^13^ (containing a 4-1BB costimulatory domain) and included a truncated CD34 reporter as a transduction marker (**Extended Data Fig. 1a**). We optimized the timing and dosing of the MM.1S myeloma model such that unedited BCMA CAR T cells and edited BCMA Mario-CAR T cells showed equivalent anti-tumor activity, and relapse began 21 days after CAR T cell transfer (**Extended Data Fig. 1b**).

Cytokine conditions during *ex-vivo* T cell product manufacturing have significant effects on T cell phenotype and proliferation^14–18^. To understand how *in vitro* culture conditions may alter the expansion and persistence of different knockout T cells *in vitro* and *in vivo*, we generated BCMA Mario-CAR T cells cultured in IL-2 or a combination of IL-7 and IL-15, both of which are commonly used to manufacture CAR T cells (**Fig. 1b**). Healthy human T cells from three normal donors were activated with anti-CD3/CD28 beads, cultured in the specified cytokine(s) throughout, and transduced with the Mario and CAR lentiviral vectors. A “baseline” sample of cells was collected 48 hours after lentiviral transduction, as a measure of library representation in cells prior to genome-editing. Following continued expansion in cytokines for 4 additional days, T cells were electroporated with Cas9 mRNA, cultured for 48 hours to enable genome editing and reduction in surface CD3 following editing of the *TRAC* locus, and subsequently enriched through CD3-negative selection (**Fig. 1b and Extended Data Fig. 1c**). We used flow cytometry to characterize the efficiency of CRISPR-editing in BCMA CAR T cells, hereafter “Mario-CAR T cells,” and determined that the anti-BCMA CAR was expressed on ∼41-58% of the CRISPR-edited, CD3-negative T cell population (**Extended Data Fig. 1d**). The CD3-negative enriched T cells were expanded an additional 5 days in presence of cytokines prior to transfer into mice bearing BCMA-positive MM.1S myeloma tumors. A sample of cells from each donor was collected at the “End of Production (EOP)” to assess the effects of gene deletion on *in vitro* expansion of T cells (**Fig. 1b**). To evaluate the effects of genetic perturbations on CAR T cell expansion and persistence *in vivo*, mice were sacrificed either 7 (“early in vivo”) or 21 days (“late in vivo”) after CAR-T injection, and cells were isolated and enriched via NGFR-positive selection from bone marrow (**Fig. 1b**).

We isolated genomic DNA from the collected *in vitro* and *in vivo* samples for PCR amplification and sequencing of the integrated sgRNAs^19^. Replicate screens were conducted using Mario-CAR T cells generated from multiple healthy human donors. All screens showed excellent replicate correlation and sgRNA recovery, as both gene-targeting and control sgRNAs formed a normal distribution across all timepoints and donors, indicating that our sgRNA level representation in the *in vivo* screen was sufficient for rigorous hit calling for both enriched and depleted genes (**Extended Data Fig. 1e-g**). Additionally, there was a high correlation of enriched and depleted sgRNAs between donors both *in vitro* and *in vivo* **(Extended Data Fig. 1h)**.

We evaluated the enrichment and depletion of sgRNAs in Mario-CAR T cells generated through either expansion protocol over time in several contexts: 1) after 11 days of *in vitro* culture and cytokine-mediated expansion compared to original library representation in unedited cells (“End of Production vs. baseline”); 2) at 7 days *in vivo* compared to the day of injection (“Early in vivo vs. End of Production”); and 3) after 21 days *in vivo* compared to the day of injection (“Late in vivo vs. End of Production”) (**Fig. 1c, Extended Data Fig. 2a**). In EOP Mario-CAR T cells, we observed strong depletion of sgRNAs targeting known common essential genes such as *DNMT1*, *PCBP2* and *SMARCB1*, as well as the key T cell transcriptional regulator *IRF4*. We also observed striking enrichment of sgRNAs targeting *RASA2* (log-fold change (LFC): 2.89; p<0.0001), consistent with previous reports that *RASA2* ablation enhances T cell proliferation *in vitro*^12^, enrichment of sgRNAs targeting *SOCS1*, a negative regulator of JAK1^20,21^, and, uniquely in the Mario-CAR T cells generated through IL-2 expansion, depletion of sgRNAs targeting *IL2RA* (LFC: –7.23; p<0.0001), consistent with its requirement for IL-2 driven T cell proliferation *in vitro* (**Fig. 1c, Extended Data** Fig 2a). Collectively, these *in vitro* observations indicated that our sgRNA library delivery and CRISPR editing was robust and could recover known biology.

To understand which genetic modifications specifically enhance T cell persistence *in vivo*, we next compared the abundance of sgRNAs at early or late *in vivo* time points to the library representation in EOP Mario-CAR. We observed that loss of the common essential genes *NDUFB10* and *ELOB* led to strong depletion both early *in vivo* and late *in vivo*. In the early *in vivo* condition, the most highly enriched sgRNAs targeted *PTPN2* (LFC: 2.41; p<0.0001), a negative regulator of JAK/STAT and TCR signaling (**Fig. 1c**)^22,23^. Deletion of the RNA regulatory genes *ZC3H12A* (REGNASE-1) and *RC3H1* (ROQUIN-1), known to play important roles in T cell responses^24–28^, also resulted in enhanced T cell expansion early *in vivo* (**Fig. 1c**). As long-term expansion and persistence is a significant challenge in CAR-T therapeutic efficacy, we next assessed sgRNA enrichment late *in vivo*, and observed that *CDKN1B* KO T cells were the most abundant compared to cells at injection (LFC: 3.49; p<0.0001). We also observed a significant enrichment of sgRNAs targeting either *SOCS1* or *PTPN2* at the later time point *in vivo,* suggesting a long-term beneficial effect of increased JAK/STAT activation, as well as enrichment of cells lacking TGFBR, a signaling pathway that has been previously linked to enhanced CAR-T persistence^29^.

To directly compare gene effects across the life cycle of Mario-CAR T, we juxtaposed enrichment of sgRNAs targeting the top gene for each of the previous comparisons and observed that despite strong enrichment at EOP, sgRNAs targeting *RASA2* conferred no benefit to Mario-CAR T expansion *in vivo* (**Fig.1d**). In contrast, sgRNAs targeting *PTPN2* were enriched in both *in vivo* time points despite no discernible effect during *in vitro* culture, while sgRNAs targeting *CDKN1B* showed modest enrichment during *in vitro* expansion and the largest enrichment after 21 days *in vivo* (**Fig. 1d**). Although there were some differences in depleted sgRNAs depending on the cytokines used during *in vitro* production, notably *IL2RA* and *DNMT1*, the overall depletion and enrichment of sgRNAs were largely concordant between manufacturing approaches (**Fig. 1e**). We set out to more comprehensively characterize patterns of enrichment and depletion of different guides over time *in vitro* and *in vivo*, by examining z-scored guide abundance at each time point for each cytokine expansion protocol (**Fig. 1f, Extended Data** Figure 2b). As might be expected, the impact of loss of *IRF4* or *STAT3*, critical mediators of TCR-activated proliferation and differentiation, during *in vitro* production led to durably low abundance *in vivo*. Cells lacking *IL2RA* showed reduced survival *in vivo* regardless of the differential sensitivity to its loss during in vitro expansion in IL-2 or IL-7/15 (**Fig. 1f, Extended Data** Figure 2b).

While loss of RASA2 led to enhanced proliferation *in vitro*, leading to increased abundance at EOP, these cells exhibited diminished abundance at both time points *in vivo*. In contrast, cells lacking *PTPN2* or *CDKN1B* showed a progressive increase in frequency *in vitro* and *in vivo*, regardless of the cytokines used during *in vitro* production (**Fig. 1f, Extended Data** Figure 2b). Collectively, our screening data revealed that many genes play variable roles regulating CAR T expansion *in vitro* and *in vivo* over time, and suggest that *in vitro* models may be insufficient for identification of genes important for *in vivo* persistence.

## Perturb-seq identifies features associated with enhanced persistence *in vivo*

To understand how various gene deletions affect T cell transcriptional state and frequency after 21 days *in vivo*, we selected a subset of the top enriched and depleted genes from our original screens (*CDKN1B*, *IL2RA*, *PTPN2*, *RASA2*, *RC3H1*, *SOCS1*, *TGFBR2*, *ZC3H12A,* and *CD160* as a gene-targeting negative control) to evaluate by Perturb-seq **(Extended Data Table 4)**. We designed 4 sgRNAs targeting each gene, as well as 16 intergenic control sgRNAs, for a total library size of 52 sgRNAs. Guide RNAs were cloned into the original screening vector that included a paired, fixed *TRAC* sgRNA to enable CD3-negative selection of CRISPR-edited T cells. Following transduction with the vectors containing the sgRNA library and the CAR, engineered T cells were enriched by negative magnetic bead selection and transferred into mice previously engrafted with MM.1S myeloma (**Fig. 2a**). After 21 days, mice were euthanized and the modified CAR-T cells were isolated from the bone marrow by NGFR-positive selection for analysis by droplet-based scRNAseq (see Methods). Leiden clustering of 18,680 cells generated 11 cell clusters (**Fig. 2b**). Clustering was primarily driven by transcripts associated with lineage, cell cycle, and transcriptional states including exhaustion, effector, and memory **(Fig. 2c-e, Extended Data Fig. 2c, Extended Data Table 5)**. To link the single cell transcriptional states with gene loss-of-function, we focused our analysis on a subset of 4,991 cells in which we detected a single genetic perturbation per cell (obligate pair of gene– and *TRAC*-targeting sgRNA). We examined the distribution of detected perturbations across clusters, compared to cells containing an intergenic control sgRNA and those with no detected sgRNA (no guide) cells (**Fig. 2f and Extended Data Fig. 2d**). We observed that the majority of no guide and intergenic KO cells were represented in the early effector CD8^+^, late effector CD8^+^, and memory-like CD4^+^ clusters. We excluded *RASA2*, *CD160*, and *IL2RA* from our analysis due to poor cell recovery (<100 cells each). Cells lacking *PTPN2*, *RC3H1* and *ZC3H12A* enriched amongst proliferating S phase Effector CD8^+^ and proliferative CD8^+^ cells (**Fig. 2f**). *CDKN1B* KO T cells were specifically enriched in the proliferative CD8^+^ cluster, while *TGFBR2* KO cells specifically enriched within the progenitor exhausted cluster (**Fig. 2f**). *SOCS1* and *RC3H1* KO caused an enrichment of proliferative CD4^+^ T cells (**Fig. 2f**). Thus, deletion of genes that promote CAR T cell persistence *in vivo* have differing effects on the transcriptional profile of CAR T cells in the tumor microenvironment.

**Figure 2.**
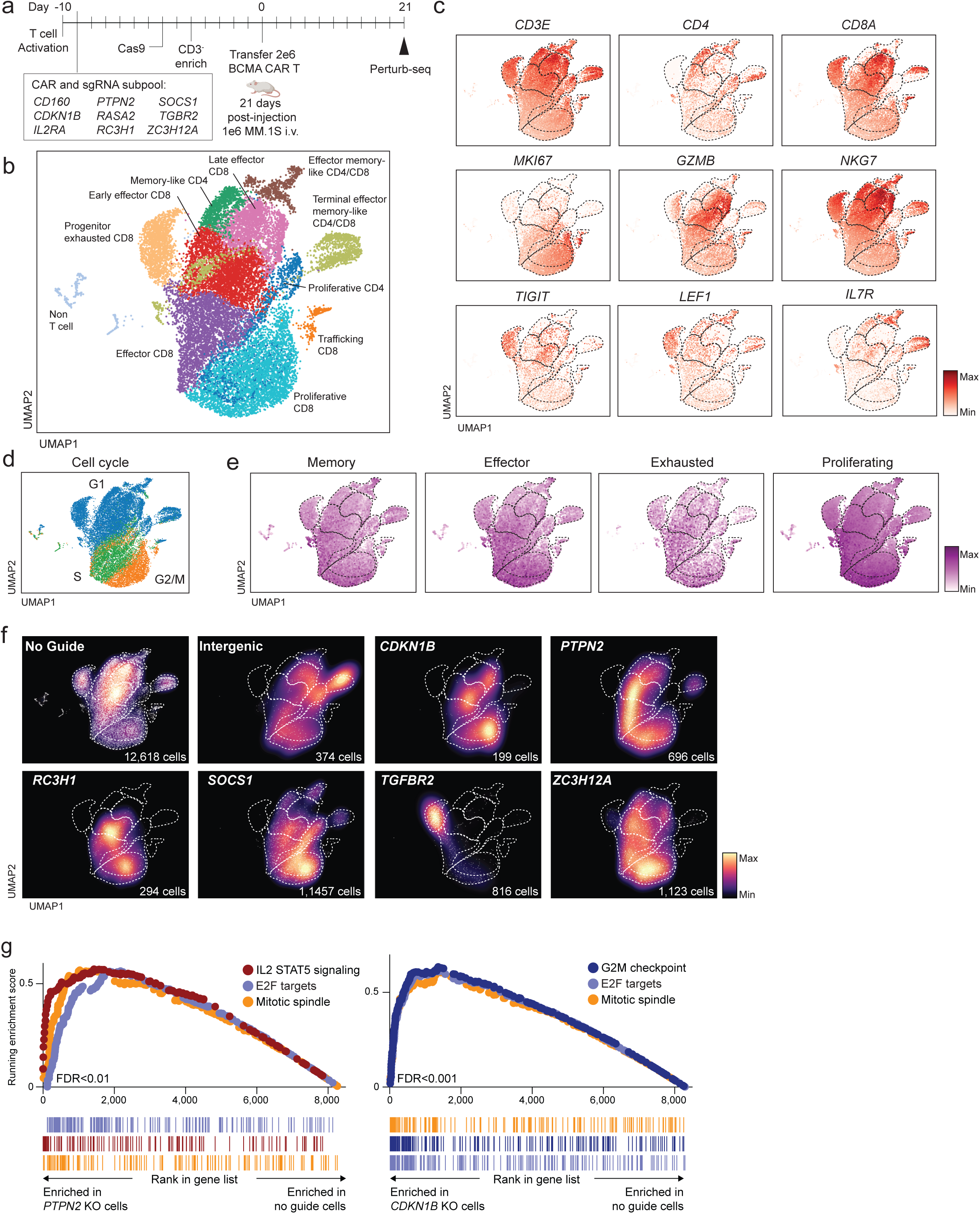
*In vivo* Perturb-seq characterization of BCMA CAR T cells. **a.** Diagram of the perturb-seq workflow. Cells were prepared as described in detail in **Fig. 1b**. The perturb-seq library featured a selected fraction of the Mario library genes (intergenic controls, *CD160*, *CDKN1B*, *IL2RA*, *PTPN2*, *RASA2*, *RC3H1*, *SOSC1*, *TGBR2*, *ZC3H12A*). Modified CAR T cells (2E6 double positive cells) were transferred into NSG mice bearing MM.1S multiple myeloma. After 21 days, the CAR T cells were isolated from bone marrow for droplet-based scRNAseq with sgRNA capture. Human T cells were from a single healthy donor (ND106) and were isolated from n=4 individual mice for scRNAseq analysis. **b.** Uniform manifold approximation and projection (UMAP) of 18,680 cells and 11 clusters identified among NGFR-enriched BCMA CAR T cells. **c.** UMAPs of expression for representative T cell phenotypic marker genes **d.** UMAPs of cell cycle score **e.** UMAPs of T cell phenotypic gene signatures **f.** Cell density projections by gene target **g.** Hallmark gene set enrichment analysis (GSEA) of pseudo-bulk pooled *PTPN2* KO (left panel) or *CDKN1B* KO (right panel) CAR T cells compared to no guide CAR T cells.

We performed gene set enrichment analysis (GSEA) on pseudo-bulk pooled knockout cells to look more in depth at knockout specific gene signatures distributed across various clusters compared to unperturbed cells (**Fig. 2g and Extended Data Table 6**). *PTPN2* KO cells showed an enrichment of the IL2 STAT5 Signaling and IL6 JAK STAT3 gene signatures, consistent with the role of PTPN2 as a dampener of JAK-STAT signaling. Cells lacking CDKN1B, PTPN2, RC3H1, SOCS1, or ZC3H12A showed significant enrichment of the hallmark G2M Checkpoint, E2F Targets, and Mitotic Spindle gene sets, suggesting that loss of function in these genes was associated with increased cell proliferation.

## *CDKN1B* KO CAR T cells display higher expansion and reduced exhaustion in *in vitro* co-culture conditions

To better understand which genetic perturbations will generate CAR T cells with favorable *in vitro* expansion and *in vivo* persistence, we performed deeper characterization of the function of CAR T cells following deletion of individual genes. We chose to focus on four genes based on their *in vitro* and *in vivo* pooled screen trajectories and transcriptional profiles. *PTPN2* and *CDKN1B* were selected based on their *in vivo* phenotypes, *RASA2* for its robust effect on *in vitro* production, and *IL2RA* for its deleterious effects during *in vitro* expansion in IL-2.

To simplify T cell production into a single lentiviral vector, we cloned the double guide cassette into the BCMA CAR vector and included a sgRNA for the chosen candidate gene or an intergenic control sgRNA (**Extended Data Fig. 3a**). We measured the expansion rate of each individual KO CAR T cell product in culture (**Fig. 3a**). *RASA2*, *PTPN2*, and *CDKN1B* KO CARs expanded significantly more compared to the intergenic control T cells during *in vitro* culture, whereas *IL2RA* KO CARs cultured in IL2 expanded significantly less. The expansion of *IL2RA* KO T cells could be rescued by culturing in IL-7/15 rather than IL-2. Gene disruption was confirmed by next-generation sequencing (NGS, **Extended Data Fig. 3b**) for *CDKN1B*, *PTPN2*, and *RASA2*; and via flow cytometry for *IL2RA* (**Extended Data Fig. 3c-d**).

**Figure 3.**
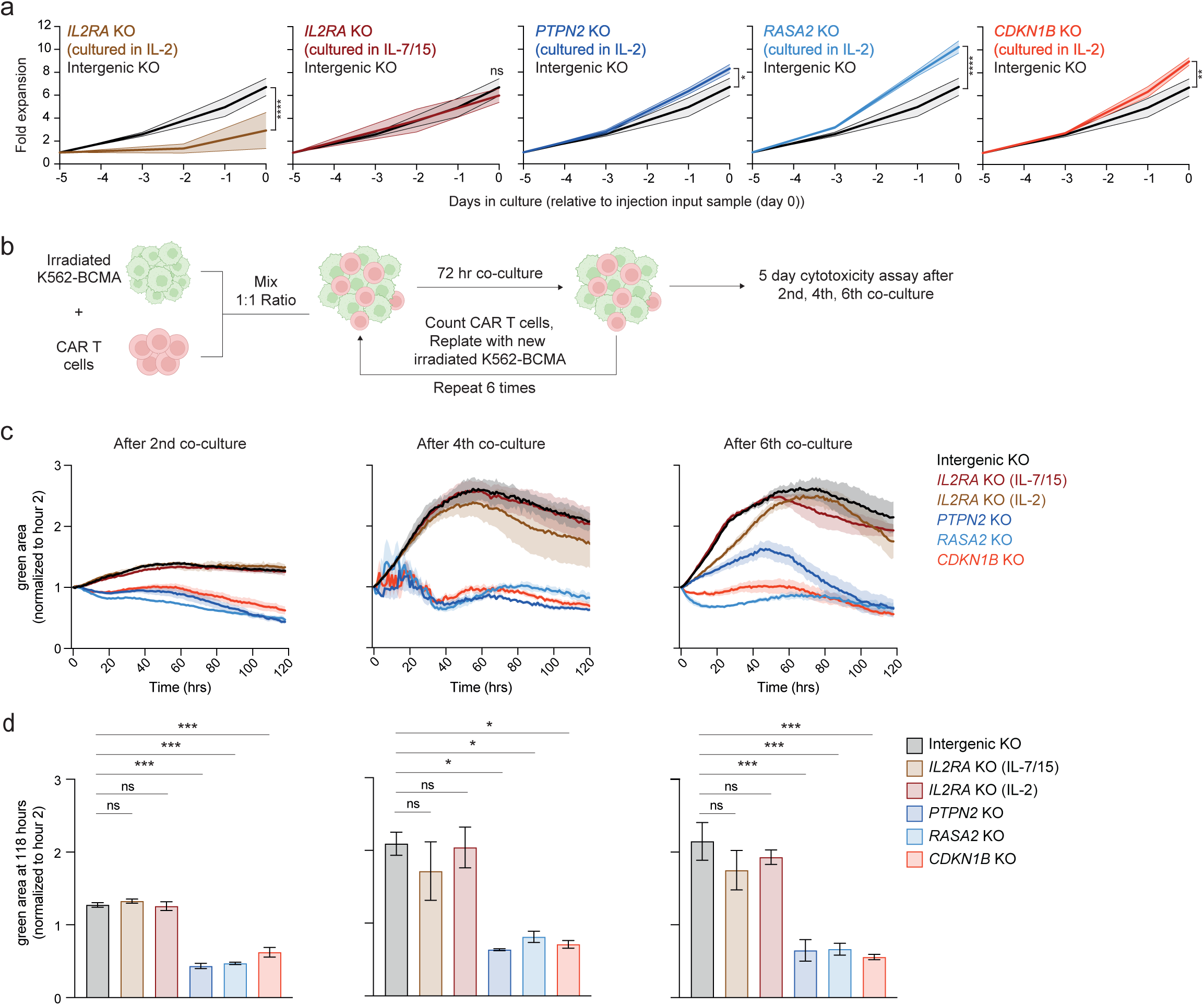
Knockout of key T cell regulators enhances BCMA CAR T cell expansion and cytotoxicity *in vitro*. **a.** Relative expansion of knockout CAR T cells during production in IL-2. Data represent one or two technical replicates from CAR T cells generated from two normal donors (ND116, ND202), measured as their fold expansion (fold change in CAR+ cells in the culture over time) *in vitro* following CD3 negative selection. Data presented as mean +/-SEM. Statistical significance was measured by two-way ANOVA with Tukey’s multiple comparison test. **b.** Schematic overview of repetitive stimulation assay. **c.** Real-time cytotoxicity assay of CAR T cells (taken from different restimulation time points (2nd, 4th, and 6th)) co-cultured with irradiated K562-BCMA target cells at a 1:1 effector:target (E:T) ratio (upper panel), with quantification of total tumor growth after 118 hours of co-culture (lower panel). Tumor cell growth is shown as the total green area relative to day 0 (tumor seeding). Data represent technical replicates from CAR T cells generated from one normal donor (ND116). Statistical significance was measured by one-way ANOVA with Tukey’s multiple comparison test. Data presented as mean +/-SEM. **b** was created with Biorender.com. *p<0.05, **p < 0.01, and ****p < 0.0001, ns: non significant.

To investigate if each individual gene KO influenced *in vitro* CAR T cell killing capacity, we performed luciferase-based killing assays at different effector to target (E:T) ratios using the human myeloma cell line MM.1S (**Extended Data Fig. 4a**) and found no significant differences. However, this assay only measured the response to acute antigen exposure. To better mimic the chronic antigen exposure CAR T cells experience *in vivo*, we performed a repetitive stimulation assay in which CAR T cells were stimulated every 72 hours with irradiated K562 cells, transduced to express BCMA, at an E:T ratio of 1:1 (**Fig. 3b**). Ablation of *CDKN1B*, *PTPN2*, or *RASA2* preserved cytotoxicity over multiple rounds of stimulation compared to intergenic KO and *IL2RA* KO CAR T cells when tested in a 5-day cytotoxicity assay with RPMI-8226 (effector to target ratio 1:1), which failed to control the expansion of tumor cells after repeated exposures (**Fig. 3c-d**).

Collectively, these data demonstrate that loss of *CDKN1B*, *PTPN2*, or *RASA2* enhances T cell cytotoxicity and prevents T cell dysfunction *in vitro* during repetitive antigen encounter.

## *CDKN1B* KO enhances CAR T cell antitumor activity and persistence in vivo

After evaluating the *in vitro* performance of our knockout CAR T cells, we sought to further characterize them *in vivo*. Following engraftment of MM.1S multiple myeloma, we treated mice with *PTPN2*, *CDKN1B*, *RASA2, IL2RA*, or intergenic control BCMA CAR T cells, all cultured in IL-2 during production. Mice that received *CDKN1B* KO-BCMA CAR T cells exhibited prolonged tumor control, while mice receiving *PTPN2* KO T cells showed only transient control of tumor growth, but later relapsed similar to mice given intergenic KO CAR T cells (**Fig 4a**). *RASA2* deficient CAR T cells had similar *in vivo* efficacy to intergenic KO CAR T cells despite enhanced cytotoxicity *in vitro*, with mice relapsing 28 days after CAR T cell injection. *IL2RA* KO T cells showed marked lack of tumor control and reduced overall survival.

**Figure 4.**
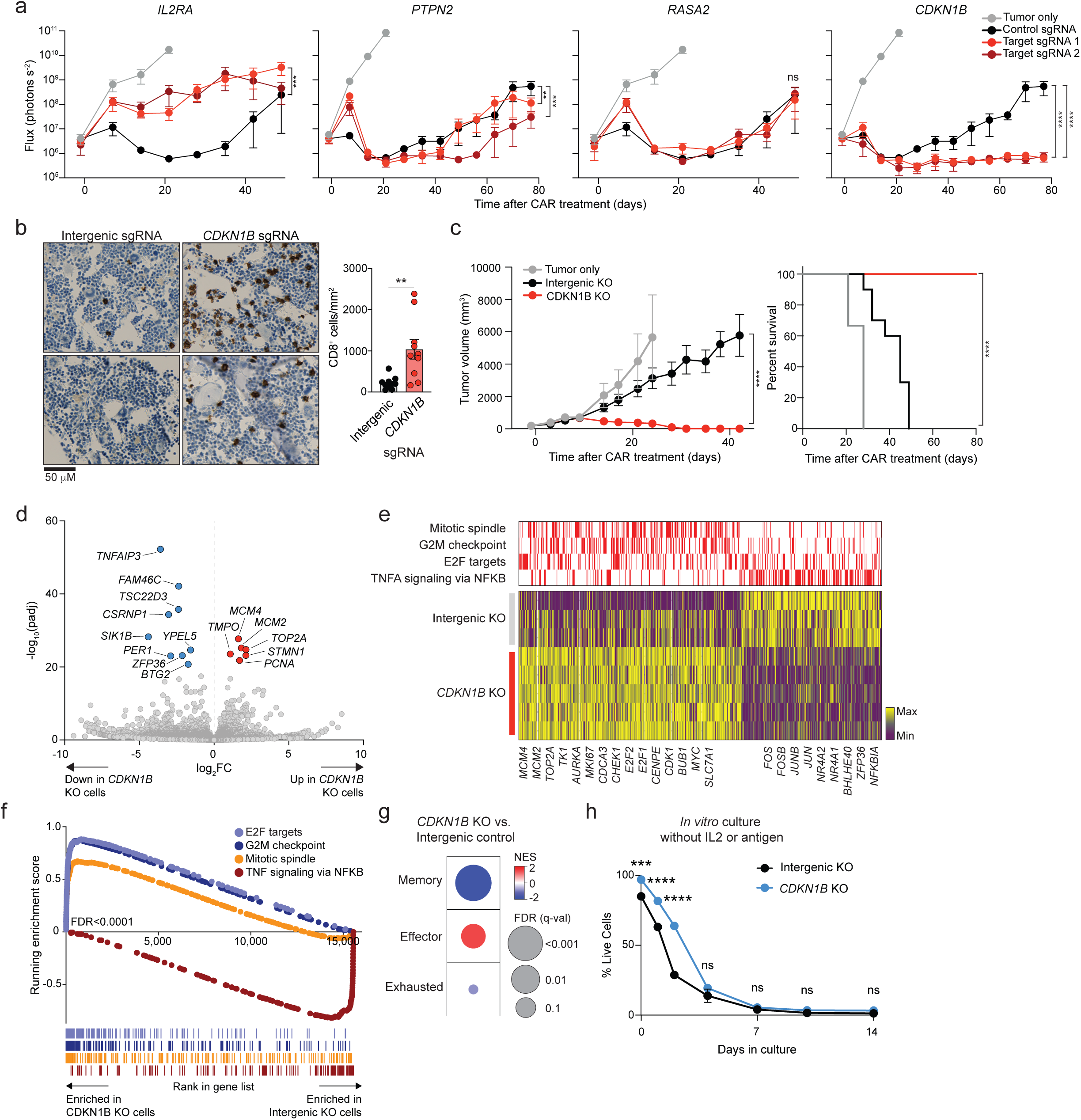
*CDKN1B* KO enhances CAR T cell function and persistence against myeloma. **a.** MM.1S tumor burden as measured by bioluminescent imaging (BLI) of mice treated with intergenic control KO, *IL2RA* KO, *PTPN2* KO, *RASA2* KO or *CDKN1B* KO BCMA CAR T cells. NSG mice were injected intravenously with 1E6 MM.1S followed by transfer of CAR T cells 21 days later. n=5 mice per group from one healthy donor (ND116). Data presented as mean +/-SEM. Statistical significance was measured compared to the intergenic KO CAR T cell treated group at day 77 (*CDKN1B* KO, *PTPN2* KO) or day 49 (*IL2RA* KO, *RASA2* KO) as measured by two-way ANOVA with Tukey’s multiple comparison test. **b.** CD8 immunohistochemistry staining of bone marrow from spine at day 21 after CAR T cell transfer (left panel). Quantification of CD8+ cells per mm^2^ from 5 independent areas of n=2 mice (right panel). Data presented as mean +/-SEM with individual data points. Statistical significance was measured using two-tailed Student’s t-test. **c**. RPMI-8226 tumor burden (left panel) and overall survival (right panel) of mice treated with intergenic control KO or *CDKN1B* KO BCMA CAR T cells. NSG mice were subcutaneously injected with 5E6 RPMI-8226 followed by CAR T cell transfer 14 days later. n=5 mice per group from two healthy donors (ND116, ND202) (10 mice total for each group). n=3 mice for tumor only group. Tumor volume was tracked by caliper measurements (mm^3^). Data presented as mean +/-SEM for tumor burden. For tumor burden, statistical significance was measured compared to the intergenic KO CAR T cell treated group at day 42 measured by two-way ANOVA with Tukey’s multiple comparison test. For overall survival, statistical significance was measured by log-rank (Mantel–Cox test) for Kaplan–Meier curves. **d.** Volcano plot of bulk RNAseq from intergenic control KO or *CDKN1B* KO BCMA CAR T cells isolated from the bone marrow of mice 21 days after transfer to NSG mice engrafted with MM.1S myeloma. n=3 mice for intergenic control and n=5 mice for *CDKN1B* KO (ND116). Select up-regulated genes in *CDKN1B* KO cells are indicated in red and down-regulated indicated in blue. **e.** Heat map showing relative expression of genes within select hallmark gene sets. Genes contained in a gene set are indicated in red and select genes labeled below**. f.** GSEA of intergenic control KO and *CDKN1B* KO CAR T cells. FDR for all was <0.0001. **g.** Heatmap of RNA-seq-derived GSEA for memory, effector and exhausted CD8+ T cell gene sets comparing intergenic control KO and *CDKN1B* KO CAR T cells. **h.** Intergenic and *CDKN1B* ko CAR T cells were evaluated for 14-day viability, as measured by live/dead staining with DAPI, in the absence of IL-2 and antigen expressing tumor cells. Statistical significance for individual days between intergenic control KO and *CDKN1B* KO CAR T cells was conducted by two-way ANOVA with Tukey’s multiple comparison test. Data presented as mean +/-SEM. **p < 0.01, ***p < 0.001, and ****p < 0.0001, ns: non significant.

Based on the *CDKN1B* KO cells enrichment in the late *in vivo* pooled screen and their striking improvement in tumor control, we decided to further characterize *CDKN1B* KO CAR T cells function *in vivo*. We generated *CKDN1B* KO CAR T cells from an additional human donor, which also displayed improved in vivo tumor control against MM.1S (**Extended Data Fig. 5a**). Collectively, mice receiving *CDKN1B* KO CAR T cells from either donor had improved overall survival compared to intergenic control KO CAR T cells (**Extended Data Fig. 5b**). We performed a total marrow harvest (femur, tibia, spine) at day 21 after T cell transfer and stained for CD8 to quantify T cell abundance in the bone marrow. Mice treated with *CDKN1B* KO CAR T cells had more CD8^+^ cells compared to those treated with intergenic KO CAR T cells (**Fig. 4b**). These data suggest that loss of *CDKN1B* increases CD8^+^ CAR T cell expansion and/or persistence in vivo.

Next, we validated the therapeutic efficacy of *CDKN1B* KO CAR T cells using a different xenograft model of multiple myeloma. Mice were engrafted subcutaneously with the RPMI-8226 myeloma cell line and treated with BCMA CAR T cells 14 days later^30^. *CDKN1B* KO CAR T cells displayed superior anti-tumor activity as well as increased survival (**Fig. 4c**).

To better understand how the loss of *CDKN1B* enhances T cell function we performed bulk RNAseq on cells isolated from bone marrow 21 days after CAR T cell transfer using the MM.1S myeloma model (**Fig. 4d,e**). Differential gene expression analysis identified many upregulated cell cycle genes in *CDKN1B* KO CARs including *MCM4*, *MCM2*, *TOP2A*, and *PCNA*. The expression of key regulators of NFκB and AP1 transcriptional activity *NFKBIA*, *JUN*, and *FOS* were lower in *CDKN1B* KO CARs. Additionally, *CDKN1B* KO CARs had lower expression of the transcription factors *ZFP36*, *NR4A1*, and *NR4A2*. GSEA amongst the hallmark gene sets showed that *CDKN1B* KO T cells were highly enriched for genes in the E2F Targets, G2M Checkpoint, Mitotic Spindle and MYC Targets V1, whereas the TNFA Signaling via NFκB signature was more enriched in intergenic control CARs **(Fig. 4f)**. We performed additional GSEA for genes differentially expressed amongst effector, memory and exhausted CD8 T cells and found that *CDKN1B* KO T cells displayed a relative enrichment of the effector gene signature and a relative depletion of the memory gene signature **(Fig. 4g)**. These data suggest that *CDKN1B* KO CARs have increased proliferation through enhanced E2F transcriptional activity. This increase in proliferation corresponds with a reduction in CAR-mediated AP1 and NFκB signaling, potentially insulating *CDKN1B* KO cells from chronic antigen exposure. We quantified cell cycle progression using a cell permeable DNA dye in vitro in the presence of BCMA^+^ tumor cells (**Extended Data Fig. 5c)**. Immediately following initial stimulation, nearly all CDKN1B KO and intergenic control cells were in G2/M phase. After 7 and 14 days, a population of intergenic control cells emerged in G0/G1, while nearly all *CDKN1B* KO cells remained in the bright G2/M population. To understand whether *CDKN1B* KO T cells had increased resistance to apoptosis or had undergone malignant transformation, we performed long term culture with and without IL2 and antigen stimulation. In the absence of antigen and IL2, CDKN1B KO CAR T cells had higher viability after 1-2 days, but this difference was lost after 7 days of culture with >95% of cells non-viable (**Fig. 4h**) These data suggest that deletion of *CDKN1B* increases BCMA CAR T cell proliferation and function without conferring any cytokine-or antigen-independent growth. In summary, our study uses a novel *in vitro-in vivo* CAR T cell screen to identify important regulators of CAR T cell expansion and persistence *in vivo*. We further demonstrate that *CDKN1B* KO BCMA CAR T cells have enhanced anti-tumor effects *in vivo* using multiple models of human multiple myeloma.

## Discussion

Current CAR T cell therapies face intrinsic and extrinsic tumor resistance mechanisms, severely reducing CAR persistence and, thus, preventing long-term remission^5,31^. Rather than ablating one gene at a time, large-scale CRISPR loss of function genetic screens offer a powerful discovery platform to efficiently reveal genetic perturbations that enhance CAR T cell function^8,10,32^. Here, we used pooled loss-of-function genetic screens to investigate a library of genes for their influence on CAR function and persistence *in vitro* and *in vivo*, showcasing a new approach for target discovery in cellular immunotherapy.

Therefore, we identified genetic perturbations *in vivo* that enhance CAR T cell expansion and function *in vivo*. The combined *in vitro-in vivo* screen allowed us to identify targets that may specifically alter T cell phenotype in one condition but not the other. For example, loss of *RASA2* led to greatly increased expansion *in vitro.* However, *RASA2* KO T cells did not score in our screen *in vivo* and had similar anti-tumor efficacy to control CARs *in vivo*. Ablation of PTPN2 showed early increased activity, in correlation with its strong *in vivo* screen scoring until day 7, but did not exhibit increased persistence or enhanced anti-tumor activity in the models of multiple myeloma that we tested.

Using our *in vivo* screen, we uncovered *CDKN1B* as a promising target for engineering persistent CAR T cells *in vivo*. Our results suggest that loss of *CDKN1B* enhances CAR T proliferation and promotes the expression of effector genes, leading to prolonged anti-tumor activity in xenograft models of human multiple myeloma. As previously described, loss of *CDKN1B* led to an upregulation of cell proliferation genes and downregulation of cell cycle inhibitors^33^. Notably, this increased proliferation did not drive these cells to exhaustion or dysfunction.

BCMA-directed CAR T cells with *CDKN1B* KO were less exhausted *in vitro* and had an increased proportion of CD8^+^ T cells compared to control BCMA CAR T cells. We also observed that *CDKN1B* KO CARs had increased anti-tumor activity despite chronic antigen exposure *in vitro* and *in vivo*. In addition to increased cell cycle activity, *CDKN1B* KO CAR T cells had reduced expression of NFκB transcriptional targets including other members of the AP1 transcription factor family. Since NFκB and AP1 are activated directly downstream of the 4-1BB CAR, we hypothesize that rapidly expanding *CDKN1B* KO CARs may experience less chronic antigen exposure, which may prevent them from becoming exhausted and dysfunctional. There have been recent reports of T cell lymphoma occurring after CAR T cell therapy (one case targeting BCMA^34^, and one case CD19^35^). We examined whether knockout of *CDKN1B* might lead to T cell transformation and found that the growth of *CDKN1B* KO and intergenic KO CAR T cells in the absence of cytokine and antigen was similar with >95% of cells being non-viable after 7 days. Additionally, no genetic alterations of *CDKN1B* have been observed in published clinical cases to date. To further protect against secondary malignancy clinical testing of CRISPR modified CAR T cells could include a drug inducible kill-switch mechanism for patient safety^36^.

While prior research on cell cycle and its regulator *CDKN1B* has provided valuable insights into the molecular mechanisms governing T cell proliferation, differentiation, and function, their influence on CAR T cells has so far not been investigated. Here, the top scoring *in vivo* gene candidate *CDKN1B* is uncovered as a promising target for enhanced persistence of BCMA-targeting CAR T cells. With its ablation, fewer CAR T cells were in a resting state – leading not only to maintained anti-tumoral activity even after several repetitive stimulations *in vitro*, but effectively clearing tumor and avoiding relapse while significantly outperforming control BCMA CAR T cell products in two different myeloma xenograft models.

One limitation of our study is the relatively small library size (135 genes) that was chosen to specifically ensure adequate engraftment and recovery of the modified T cell library *in vivo*. Additionally, the genes within the library were curated based on those predicted to play a role in T cell function. Our study provides proof-of-concept that future *in vivo* T cell screens could be possible with larger sgRNA libraries targeting additional genes, including those that may not have known functions in T cell biology.

In summary, our findings demonstrate that *CDKN1B* ablation increases the functional persistence of CAR T cell therapy in multiple myeloma, which may prolong the duration of long-term remission for patients. Furthermore, our data suggest that there are key differences in the selective pressures that occur during antigen stimulation *in vitro* compared to the chronic antigen exposure and physiological environment of *in vivo* mouse models of human cancer.

## Methods

### Study design

This study was designed to identify the loss of genes that enhance persistence and function of CAR T cells targeting BCMA for the treatment of myeloma. To validate the targets found in the loss of function *in vivo* CRISPR screen, *in vitro* and *in vivo* functional assays were performed.

As a source of T cells, anonymized human blood samples were used with approval of the Institutional Review Board (IRB) at the Massachusetts General Hospital (MGH) and declared as ‘*non-human subjects research*’. Mice used in *in vivo* experiments were randomized prior to CAR T cell treatment. All animal work was performed according to protocols approved by the MGH Institutional Animal Care and Use Committee (IACUC).

### Mice and cell lines

All *in vivo* experiments were performed in male and female mice according to MGH Institutional Animal Care and Use Committee approved protocols. NOD-SCID-γ chain −/− (NSG) mice were bred under pathogen-free conditions at the MGH Center for Cancer Research. All mice were maintained in 12:12 h light:dark cycles at 30–70% humidity and a room temperature of 21.1–24.5 °C. All cell lines (MM.1S, RPMI-8226, and K562) were obtained from the American Type Culture Collection and maintained under conditions as outlined by the supplier; cell lines were routinely tested for mycoplasma contamination (and were negative) and authenticated by STR profiling in a 3-year cycle. Cancer cell lines used for in vivo experiments were transduced to express click beetle green (CBG) luciferase and enhanced GFP (eGFP) followed by sorting on a BD FACSAria II, FACSAria Fusion or FACSymphony S6 cell sorter to obtain a 100% transduced population.

### Construction of CARs and double guide cassette

All CAR constructs contained a CD8 hinge and transmembrane domain, 4-1BB costimulatory domain, and CD3ζ signaling domain. CAR constructs were synthesized and cloned into second-generation lentiviral vectors under the regulation of a human EF-1a promoter. BCMA (bb2121) CAR T constructs used for initial screening experiments and double transduction contained a truncated CD34 gene for evaluating transduction efficiency. A double sgRNA cassette utilizing the human U6 and H1 promoters was adapted for T cell screening by adding a fixed TRAC sgRNA and golden-gate cloning compatible BsmbI sites for variable sgRNA introduction ^37^.

The validation CAR T cell constructs also contained a CD8 hinge and transmembrane domain, 4-1BB costimulatory domain, and CD3ζ signaling domain, with the fluorescent reporter mCherry for evaluating transduction efficiency. Additionally, these constructs contained the same double guide cassette human U6 and H1 promoters with fixed TRAC sgRNA and cloning site for a variable sgRNA.

### Lentivirus and CAR T cell production

Replication deficient lentivirus was produced by transfecting plasmids into HEK293T cells, after they were expanded in R10 media (RPMI + Glutamax + HEPES (ThermoFisher Scientific, Cat-no. 72400047), supplemented with 10% FBS, penicillin, and streptomycin). Supernatant was collected at 24 and 48 hours after transfection. Filtered virus was then concentrated by ultracentrifugation on the ThermoFisher Scientific Sorvall™ WX+ Ultracentrifuge and stored at –80C.

Human T cells were purified (Stem Cell Technologies, Cat-no. 15061) from anonymous human healthy donor leukopaks purchased from the Massachusetts General Hospital blood bank under an Institutional Review Board-exempt protocol. T cells were isolated using Stem Cell Technologies T cell Rosette Sep Isolation kit. To generate CAR T cells, bulk human T cells were activated on day –14 using CD3/CD28 Dynabeads (ThermoFisher Scientific, Cat-no. 40203D) at a 1:3 T cell:bead ratio cultured in R10 media and 20 IU of recombinant human IL-2 (Peprotech, Cat-no. 200-02). Cells were transduced with CAR lentivirus at a multiplicity of infection (MOI) of 5 on day –13 and expanded with media doubling and IL-2 replacement every 2 days. For production of Mario-CAR T cells, activated cells were transduced sequentially first with the BCMA CAR and then with Mario sgRNA library lentivirus (both MOI 5). On day –8 the Dynabeads were removed via magnetic separation. The next day (day –7) the cells washed 3 times in Opti-Mem. Up to 5E6 cells were then resuspended in 100 μl Opti-MEM and electroporated with 10 μg CleanCap™ Cas9 mRNA (TriLink, Cat-no. L-7206). On day –5, CD3 negative selection was performed utilizing the EasySep™ Human APC Positive Selection Kit II (Stem Cell Technologies, Cat-no. 17661) with CD3 APC antibody (Biolegend, Anti-Human Clone OKT3, Cat-no. 317318). CAR T cells were assessed for transduction efficiency via tCD34/APC (BCMA CAR/guide library) or mCherry (validation) expression on day –4 and stored in liquid nitrogen at the end of production. For some experiments, T cells were cultured in 10 ng/ml IL7 (Peprotech, Cat-no. 200-07) and 10 ng/ml IL15 (Peprotech, Cat-no. 200-15) instead of IL2 according to the following scheme: after Dynabead activation, IL-15 (10 ng/ml) and IL-7 (10 ng/ml) is added, followed by addition of IL-15 twice per week and IL-7 once per week during CAR T cell production.

For production of Mario BCMA CAR T cells used in the CRISPR screens, three healthy normal donors (ND) were used (ND216, ND99, ND106). For the IL7/15 screen, ND216 and ND106 were used. Validation experiments of the respective gene knockouts were tested in T cells of two additional healthy donors (ND116, ND202).

Individual genetically modified BCMA CAR T cells and control T cells were grown in a similar manner with the following modifications: cells were transduced once with the construct of interest, and cells were de-beaded, washed 3 times in Opti-Mem, up to 5 × 10^6^ cells were resuspended in 100 μl Opti-MEM and electroporated with 10 μg Cas9 mRNA on day –7. Untransduced T (UTD) cells from corresponding donors were grown at the same time for controls.

### Mario library curation and preparation

The ‘Mario’ sgRNA library targeted 135 genes with known or proposed functions in mouse or human T cells. 8 guides per gene were designed using the Broad Institute Genetic Perturbation Platforms CRISPICK tool. The library also included 100 sequences targeting intergenic sites as negative controls. All targeted genes and sgRNA sequences are included in Extended Data Table 1.

### *In vivo* loss of function CRISPR screen

NSG mice were injected intravenously with MM.1S tumor cells on day –21 to engraft. CAR T cells were prepared as described above and injected intravenously on day 0. Mice were euthanized at either day 7 or day 21, and femur, tibia, and spine were collected for isolation of marrow infiltrating T cells. From these samples, Mario-CAR T cells were selected using the EasySep™ Human PE Positive Selection Kit II (Stem Cell Technologies, Cat-no. 17654) with PE NGFR antibody (Biolegend, Cat-no. 345106).

After NGFR positive selection, genomic DNA was isolated using the Qiagen QIAmp DNA Mini kit (Cat-no. 51304). sgRNA sequences were PCR amplified from genomic DNA and sequenced on an Illumina MiSeq using MiSeq Reagent Kit v2 50 cycle (Cat-no. MS-102-2001). Samples were processed according to our previously published *“In vivo* CRISPR screening protocol”^19^.

### Selection of gene candidates for validation

Specific guides used to create knockout CAR T cells were the top two scoring guides from the primary screen:

*CDKN1B* guide 1 (GGAGAAGCACTGCAGAGACA) and guide 2 (GCAGTGCTTCTCCAAGTCCC), *IL2RA* guide 1 (TGTGTAGAGCCCTGTATCCC) and guide 2 (ACTGCAGGGAACCTCCACCA), *PTPN2* guide 1 (GCGCTCTGGCACCTTCTCTC) and guide 2 (GCACTACAGTGGATCACCGC), *RASA2* guide 1 (GGGTACGATAAACTTCTTCC) and guide 2 (ATGAATAGTACATACCTATA).

### Knockout confirmation by next-generation sequencing

Genomic DNA was isolated from 1E6 T cells using the QIAamp DNA Mini Kit (Qiagen, Cat-no. 51304). After PCR, next-generation sequencing was performed (complete amplicon sequencing) by the Massachusetts General Hospital DNA Core.

### Perturb-seq

The Perturb-seq pool included 4 guides per gene targeting *CD160, CDKN1B, IL2RA, PTPN2, RASA2, RC3H1, SOCS1, TGFBR2, ZC3H12A*, and 16 intergenic control sequences. All sgRNA sequences in the pool are included in Extended Data Table 4. After NGFR positive selection, we performed droplet based scRNA seq using the 10x Chromium Next GEM Single Cell 5’ Reagent Kit v2 (Dual Index) with Feature Barcode technology for CRISPR Screening. Sequencing was performed on an Illumina NovaSeq 6000 instrument.

### Bulk-RNA sequencing

T cells were isolated from the spine and femurs of intergenic control KO and *CDKN1B* KO BCMA CAR T cell treated animals 21 days after T cell transfer. CAR^+^ T cells were sorted on a Sony SH800 Cell Sorter. Following sorting, RNA was isolated using the Qiagen RNeasy Micro Kit (Cat-no. 74004). Bulk RNA sequencing libraries were prepared using the NEB Next Ultra II Directional RNA Library Prep Kit for Illumina (New England Biolabs, Cat-no. E7765) and sequenced on an Illumina NextSeq 500 instrument.

### Flow cytometry

Cells were washed with 2% FBS in PBS, incubated with antibody for 25-30 minutes at 4 C in the dark, after which they were washed twice again. For some experiments, DAPI (ThermoFisher Scientific, Cat-No. PI62247) was added to distinguish live versus dead cells prior to analysis on a BD Fortessa X20. For antibodies that required secondary staining the procedure was similar: after primary staining as described above, secondary was added (prior to addition of DAPI), stained for 20 min at 4 °C in the dark and washed twice. For flow cytometry analysis, the following antigens were stained using the indicated antibody clones: NGFR (Mouse anti-Human, Biolegend, PE Clone ME20.4, Cat-no. 345106), NGFR (Mouse anti-Human, Biolegend, APC Clone ME20.4, Cat-no. 345108), CD34 (Mouse anti-Human, Biolegend, BV650 Clone 561, Cat-no. 343623), CD3 (Mouse anti-Human, Biolegend, APC Clone OKT3, Cat-no. 317318), CD3 (Mouse anti-Human, Biolegend, BV421 Clone UCHT1, Cat-no. 562426), CD25 (Mouse anti-Human, Biolegend, APC Clone BC96, Cat-no. 302610), CD3 (Mouse anti-Human, BD Biosciences, APC-H7 Mouse Anti-Human Clone SK7, Cat-no. 641397). For the cell cycle analysis, Vybrant™ DyeCycle™ Green Stain (ThermoFisher Scientific, Cat-no. V35004) was used following the manufacturer’s instructions.

### Luciferase-based cytotoxicity assays

Assessment of cytotoxicity was performed by a co-culture of CAR T cells with CBG-expressing tumor cells (MM.1S, RPMI-8226) at the different effector to target (E:T) ratios (ranging from 10:1 to 1:100) for a time period of approximately 16 hours. A Synergy Neo2 microplate reader by Biotek was used to measure luciferase activity. Percentage specific lysis was calculated using the following formula: (target cells-only relative luminescence units (RLU) − total RLU with CAR T cells)/(target cells-only RLU) × 100%.

### Real-time cytotoxicity assay

CD9 antibody (Clone: HI9a, Biolegend, Cat-no. 312102; 4 μl in 1ml PBS) was used to coat a 48-well plate, which was incubated overnight at 4C. The next day, the plate was washed three times with PBS. RPMI-8226 tumor cells, expressing CBG-GFP, were seeded and cells were allowed to settle at 37C for 30 minutes. Next, CAR T Cells were seeded and the plates were placed in a Sartorius Incucyte live cell imager. All cells were cultured in R10 media, with images taken every 60 minutes. Analysis was performed using the Incucyte live cell imaging software.

### Repetitive stimulations

CAR T cells were co-cultured with tumor cells (irradiated BCMA expressing K562) at a 1:1 ratio (both 2.5E5). After 3 days, CAR T cells were counted by flow-cytometry and then rechallenged with fresh irradiated K562-BCMA cells in the same 1:1 ratio (2.5E5). This was repeated six times overall (six re-stimulations).

After two, four, and six re-stimulations, CAR T cells were seeded with RPMI-8226 tumor cells in the real-time cytotoxicity assay (described above) to assay CAR function after repetitive stimulation.

### *In vivo* validation models

All mouse injections were performed by one animal technician and monitoring was blinded to expected outcomes. Mouse experiments included at least 3 mice per group, with the exact numbers used for each experiment specified in the figure legend. Mice were randomized on day –1, therefore post-tumor injection and one day prior to treatment. Multiple myeloma MM.1S cells were administered intravenously with 1E6 cells in 100 μl PBS and engrafted for 21 days prior to treatment with 2E6 BCMA CAR T cells, also in 100 μl PBS given intravenously.

Mice were monitored for bioluminescent emission biweekly as previously described and euthanized as per the experimental protocol or when they met pre-specified endpoints defined by the IACUC. Aura software was used to analyze bioluminescent images.

### Pooled CRISPR Screening analysis

Guide sequences were demultiplexed and quantified using PoolQ v.2.2.0. Barcode count data was initially processed and quality checked. Samples with a ‘Normalized Match’ value < 10 were excluded from the analysis to ensure data reliability. The PCR replicate counts were summed together and biological replicates of mice were averaged together for each condition. The guide counts data was then normalized to reads per million (RPM) and log2 transformed with a pseudocount of 1. The guide distribution in the library was visualized as the density of log2rpm values. Pearson’s correlations were calculated for the library distribution in one biological replicate versus any other replicate, two averaged replicates versus any other two, and so on. The mean of all possible combinations was plotted. Z-score normalization was applied to the log2rpm data for all sgRNAs using the control sgRNA distribution as the baseline. Natural cubic splines with 4 degrees of freedom were fitted to the zlog2rpm data for each sample-control pair to calculate residuals. The z-log fold changes (zLFC) were calculated as the difference between the sample and control zlog2rpm values, while the residuals (zresid) were calculated as the deviations from the spline fit. Density scatter plots were produced, where each point represents the zlog2rpm values for a given guide in the control (x-axis) and sample (y-axis) where deviations from the spline fit line represent the residuals used in downstream analysis. Density distribution plots were generated to visualize the distribution of guide residuals across different donors and conditions. To further assess donor concordance within each screen, Pearson correlation values of log2RPM values were calculated. The correlations of log2rpm and zresid values were additionally visualized using scatter plots. These analyses confirmed that the donors were concordant. Consequently, further analysis was conducted using the averaged data from all donors.

Parallel guide abundance plots visualize log2rpm values over time for each guide associated with selected genes of interest to investigate the performance of each guide. The zresid values were plotted in a sweissogram, where each guide is represented as a red line on the distribution of all values in the given comparison. In the volcano plots, the LFC values for each gene were calculated by averaging the zresid values of the top performing guides for each condition comparison. Then the p-values are calculated by using the hypergeometric distribution to assess the significance of enrichment or depletion of a gene’s signal which are then –log10 transformed. To compare the hits in each screen the zLFC values from each screen are plotted against each other for a given comparison in a scatter plot.

### Perturb-seq analysis

Alignment and count aggregation were performed using CellRanger (v.7.1.0). Gene expression and sgRNA reads were aligned using cellranger count with default settings. Gene expression reads were aligned to the “refdata-gex-GRCh38-2020-A” human transcriptome. sgRNA reads were aligned to the library using the pattern TTCCAGCATAGCTCTTAAAC(BC). Before count aggregation and read depth normalization, there were an estimated 23,424 cells recovered across 4 replicates of day 21 samples, from one donor. Counts were then aggregated using cellranger aggr with default settings. Next, a series of quality control checks were performed using Scanpy (v.1.9.5). Cells with greater than 10% mitochondrial gene content were removed. Cells with reads less than 2,000 counts and greater than 18,000 counts were removed to ensure the overall read depth per cell is within a reasonable range. Cells that had less than 1,150 genes were excluded due to poor gene capture. To remove any remaining doublets, the Scrublet tool was employed with an expected doublet rate of 10% and 50 principal components. The resulting population of cells after standard quality control filters was 18,680 cells.

If at least one sgRNA read is detected in a cell, that cell is assigned as containing the associated guide accordingly. This resulted in a distribution of guides in cells where the majority of cells contain 1 guide, and the maximum number of guides in any given cell is 5. The data was not filtered on CRISPR read counts as there was poor guide capture and the majority of cells had 1 CRISPR read count.

Principal component analysis (PCA) and nearest neighbor graphs were calculated on a set of 10,000 highly variable genes using log-transformed gene expression data to visualize the cells on a UMAP plot. Harmony batch correction was then used to correct principal component analysis (PCA) embeddings for technical batch effects between the samples.

The cells were then grouped into 11 clusters using the Leiden algorithm with a resolution of 0.4. Leiden clusters were classified on the basis of the built-in scanpy function one-versus-rest differential expression, expression of marker genes of interest, CD4+ versus CD8+ expression, and cell cycle scoring to determine T cell subset identity. Cell cycle scoring was calculated using Scanpy with a cell cycle gene list from a previously published scRNAseq study^38^. After cells were phenotyped, we projected the distribution of cells with specific guides across clusters. This analysis highlighted which guides were more likely to be associated with certain T-cell subsets. Notably, cells containing the RASA2, IL2RA, and CD160 guides were excluded from the chi square analysis and further analysis due to low cell counts (< 100).

For each subset of guide containing cells, irrespective of T-cell identity, a pseudo-bulk expression profile was created by summing the counts of all cells containing a given guide. This data was then converted into a counts table where genes with low expression (counts<10) and TCR genes were filtered out. Subsequently, a differential expression analysis was performed to identify differentially expressed genes between each guide containing subset and the subset of cells with no guide. Ranked lists of differentially expressed genes were created using the log2foldchange values calculated by DeSeq2 (pydeseq2 v.0.4.8). These ranked lists were passed to GSEA Pre-rank to search for enriched hallmark gene sets using gseapy (v.1.0.5).

### Bulk RNA-seq Analysis

Reads were adapter– and quality-trimmed using Trimmomatic (v.0.36). Trimmed reads were quantified by pseudoalignment to GRCh38 using Kallisto (v.0.44.0). Abundance estimates were then transformed into gene counts using Tximport (v.1.8.0). Differentially expressed genes were qualified using DESeq2 (v.1.38.3) and GSEA pre-rank was performed using gseapy (v.1.0.5) to determine enriched hallmark genesets. From the Human MSigDB v.2023.2.Hs collection, the memory signature was defined as the overlapping genes between GSE9650_EFFECTOR_VS_MEMORY_CD8_TCELL_DN and GSE9650_EXHAUSTED_VS_MEMORY_CD8_TCELL_DN. The effector gene signature was defined as the overlap between GSE9650_EFFECTOR_VS_EXHAUSTED_CD8_TCELL_UP and GSE9650_EFFECTOR_VS_MEMORY_CD8_TCELL_UP. The exhausted signature was defined as the overlap between GSE9650_EFFECTOR_VS_EXHAUSTED_CD8_TCELL_DN and GSE9650_EXHAUSTED_VS_MEMORY_CD8_TCELL_UP.

### Statistical methods

Analyses were performed with GraphPad Prism 10 (version 10.2.3). Unless otherwise stated, data were presented as mean ± SEM and a two-tailed Student’s t-test or one-or two-way ANOVA tests were used. All tests were two-sided unless otherwise specified. Significance was considered for p < 0.05 as the following: *p < 0.05, **p < 0.01, ***p < 0.001, and ****p < 0.0001. For experiments with multiple groups, multiple comparisons corrections were used as indicated in the figure legends.

### Data availability

Transcriptomic data will be made available through GEO or dbGaP pending manuscript acceptance. Accession codes will be provided upon acceptance.

### Critical reagents and materials to this study

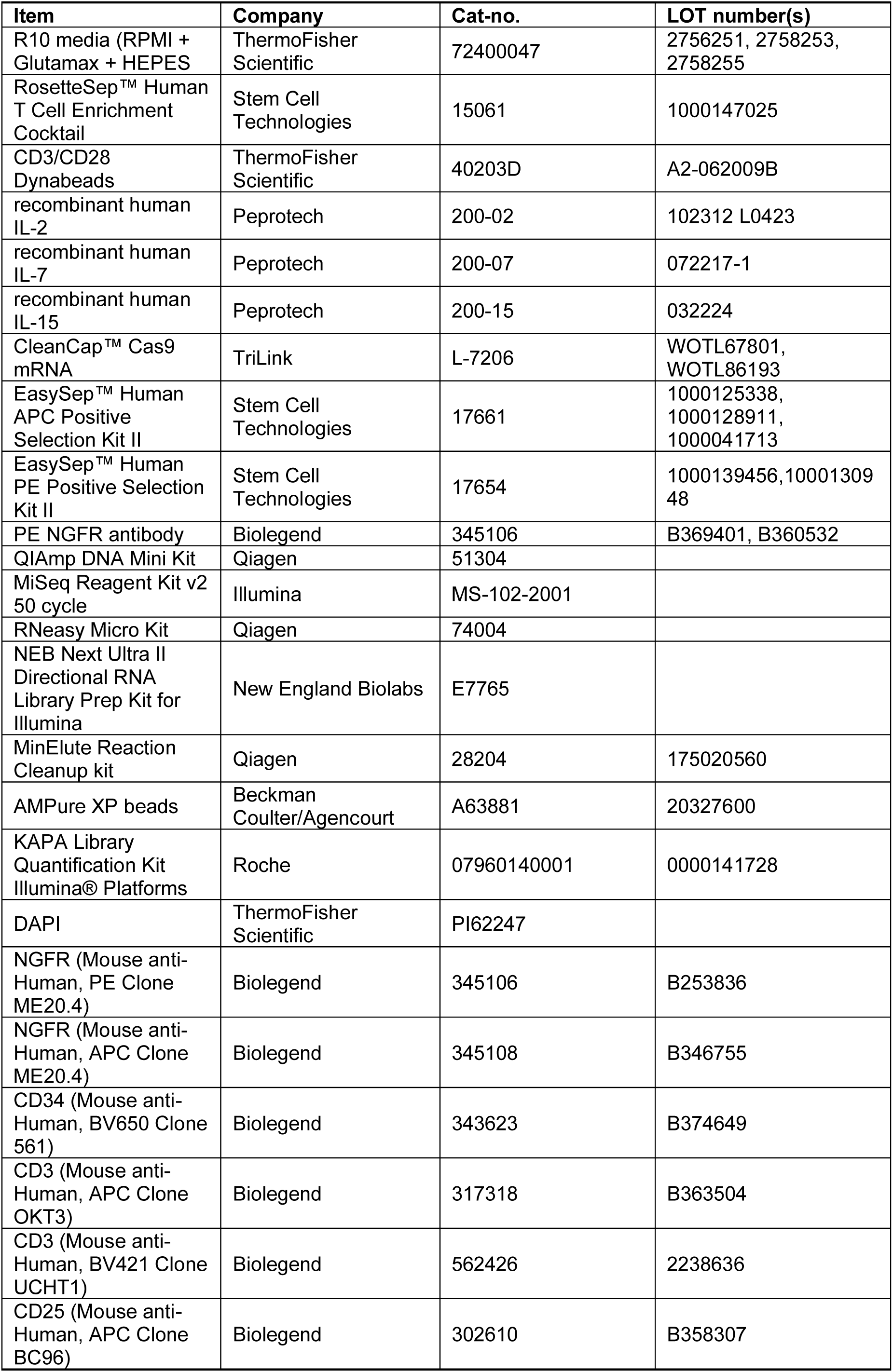
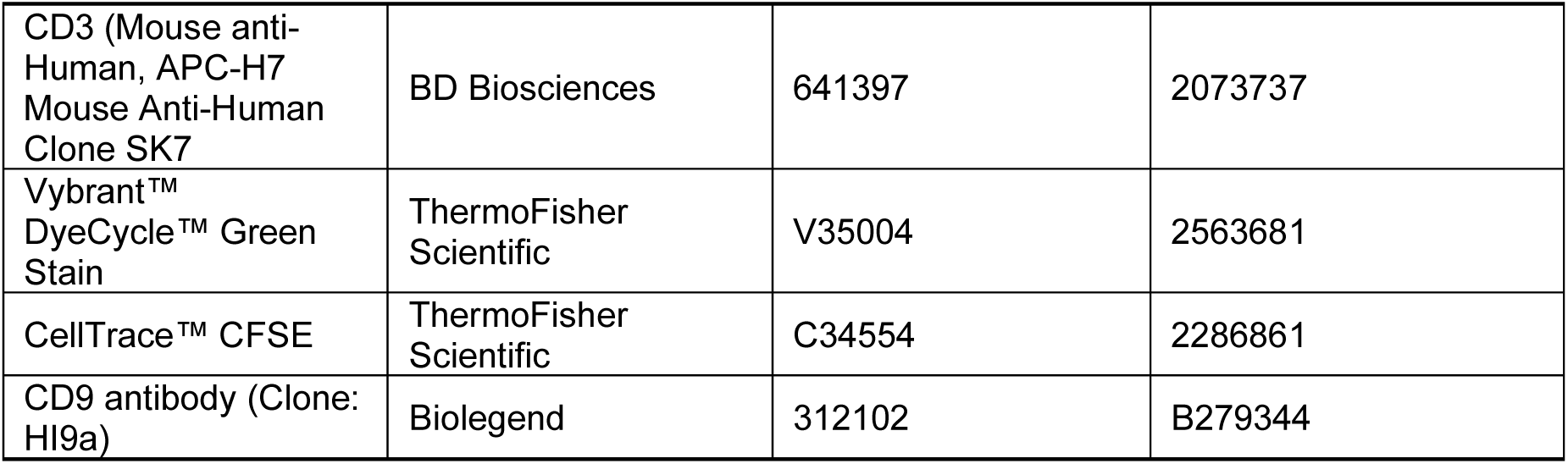

## Acknowledgments

FK received funding from the German Research Foundation (DFG; 466535590). DSB received funding from the CRIS Foundation Out-Back Fellowship Programme (outback2021_6) and from the Spanish Society of Medical Oncology (SEOM). FB received funding from the American-Italian Cancer Foundation (AICF) and the Italian Association for Cancer Research (AIRC). This work was funded by NIH R01 CA238268 (MVM) and the Krantz Breakthrough Award.

## Author contributions

Conception and experimental design: F.K., N.H.K., T.K., R.C.L., J.G.D., D.S., K.B.Y., R.T.M., M.V.M. Experimental and computational data generation and acquisition: F.K., N.H.K., T.K., G.E., C.N., S.A., A.Y.C., M.Z., A.B., M.C.K., S.G., H.W.P., M.Pezeshki, A.R., J.S.M.T.S., M.Phillips, S.P., D.S.B., E.P.D. Analysis and interpretation of data: F.K., N.H.K., T.K., G.E., C.N., S.A., A.Y.C., M.Z., A.B., M.C.K., S.G., H.W.P., M.Pezeshki, A.R., J.S.M.T.S., M.Phillips, F.B., M.B.L., R.C.L., J.G.D., D.S., K.B.Y., R.T.M., M.V.M. Manuscript writing and revision: F.K., N.H.K., C.N., T.R.B, K.B.Y., R.T.M., M.V.M.

## Corresponding authors

Correspondence to Robert T. Manguso or Marcela V. Maus.

**Extended Data Figure 1.**
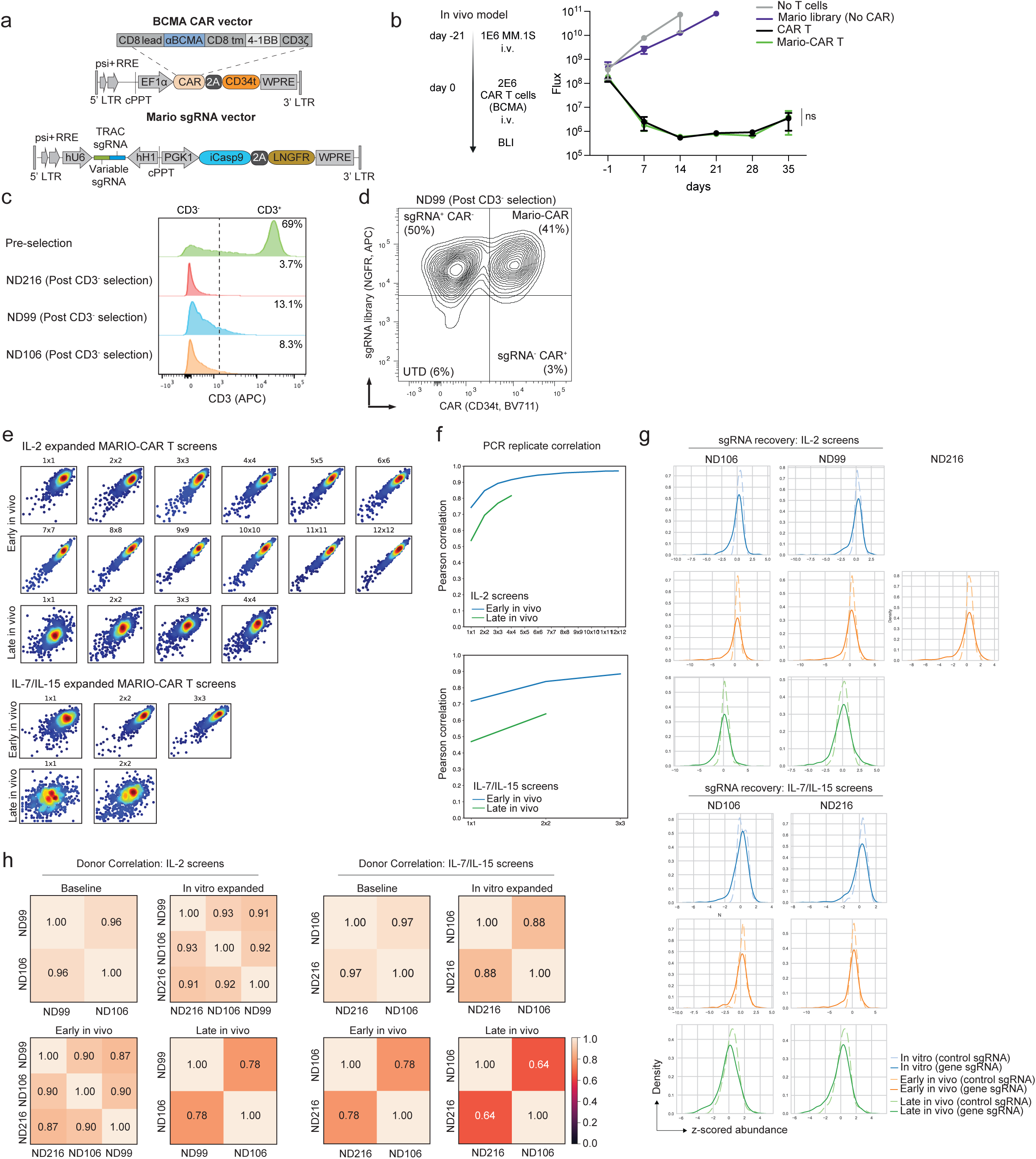
*In vivo* human CAR T cell screening results are comparable across multiple healthy human donors. **a.** Construct designs for the 4-1BB BCMA CAR (pCAR) and double-guide cassette (pGuide) containing the Mario sgRNA library (variable sgRNA). **b.** Timeline of the MM.1S stress model with 21 day tumor engraftment and 2E6 CAR treatment. Tumor growth was tracked by BLI. Data presented as mean +/-SEM. n=3 mice (tumor only, Mario library (no CAR)) and n=5 (CAR T, Mario-CAR T) per group from one healthy donor (ND202). Statistical significance amongst groups at day 35 as measured by two-way ANOVA with Tukey’s multiple comparison. **c.** CD3 expression after day –5 CD3 negative enrichment (ND216, ND99, ND106). **d.** Representative Mario-CAR T cell staining for CD34 indicating CAR transduced cells and NGFR for sgRNA library transduced cells from ND216. **e**. Replicate autocorrelation analysis scatter plots. Pearson’s correlations are calculated for the library distribution of one animal versus any other animal, two averaged animals versus any other two, and so on. The mean of all possible combinations is plotted. **f.** Quantification of replicate autocorrelation analysis. Pearson’s correlations are calculated for the library distribution of one animal versus any other animal, two averaged animals versus any other two, and so on. The mean of all possible combinations is plotted. **g.** z-scored abundance histograms of gene targeting or intergenic control sgRNAs across screening time points and conditions. **h.** Donor Pearson correlations across screening time points and conditions. ns = non significant.

**Extended Data Figure 2.**
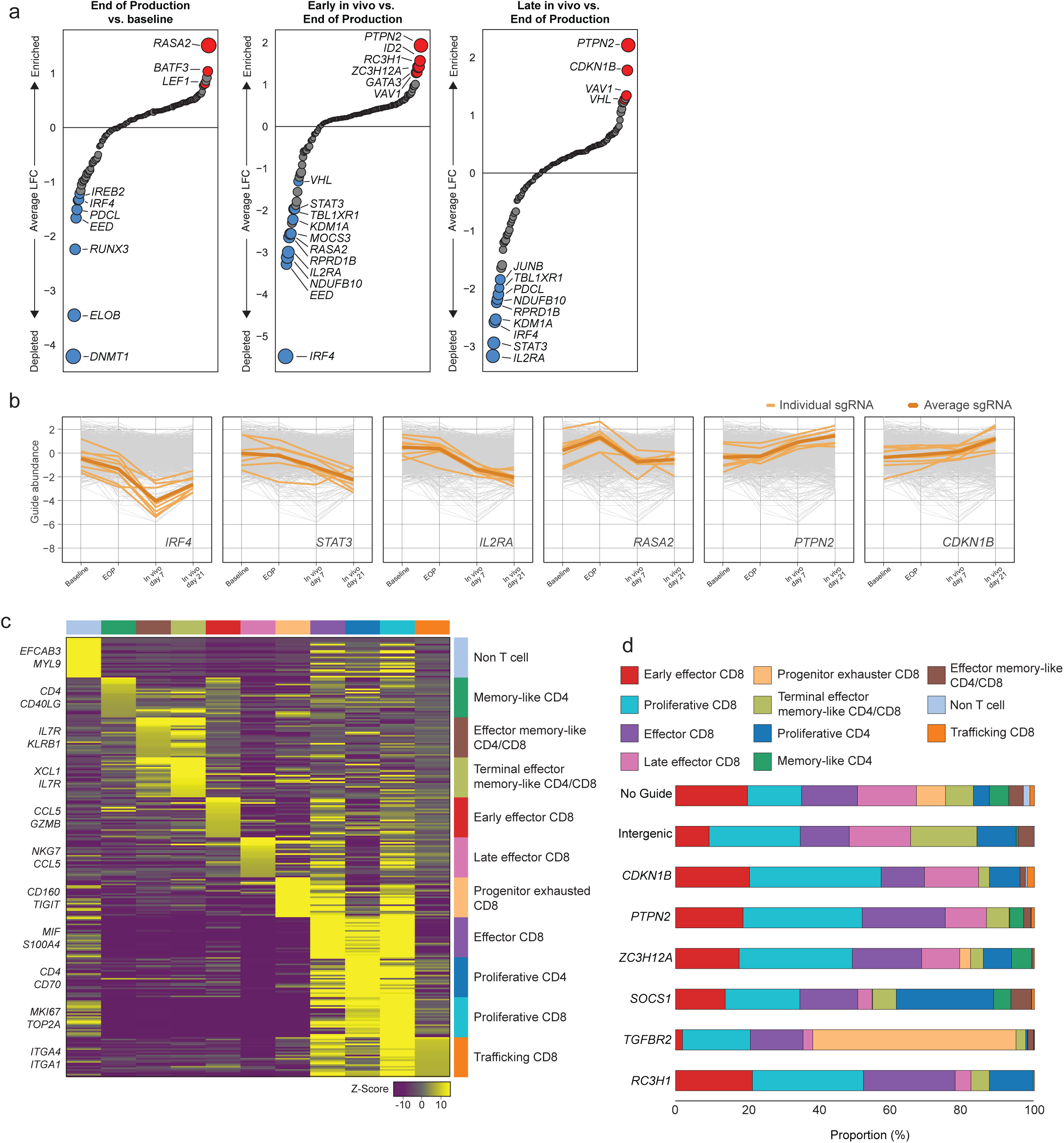
In vivo screens identify genes that modify CAR T cell abundance and transcriptional phenotype. **a.** Genes ranked by Log2(fold change) during in vitro manufacturing in IL-7/IL-15 (end-of-production vs. baseline, left panel) and early in vivo (day 7 vs. end-of-production, middle panel) or late in vivo (day 21 vs. end-of-production, right panel). Enriched genes are shown in red and depleted genes are shown in blue with circle size corresponding to –log10 (FDR). n = 2 healthy donor T cells (ND106, ND216). **b.** Abundance of sgRNAs targeting individual genes across the entire screen workflow for Mario-CAR T cells produced in IL-7/IL-15. **c.** Heatmap showing relative expression of top differentially expressed genes amongst clusters of BCMA CAR T cells. **d.** Proportions of each knockout cell type amongst clusters.

**Extended Data Figure 3.**
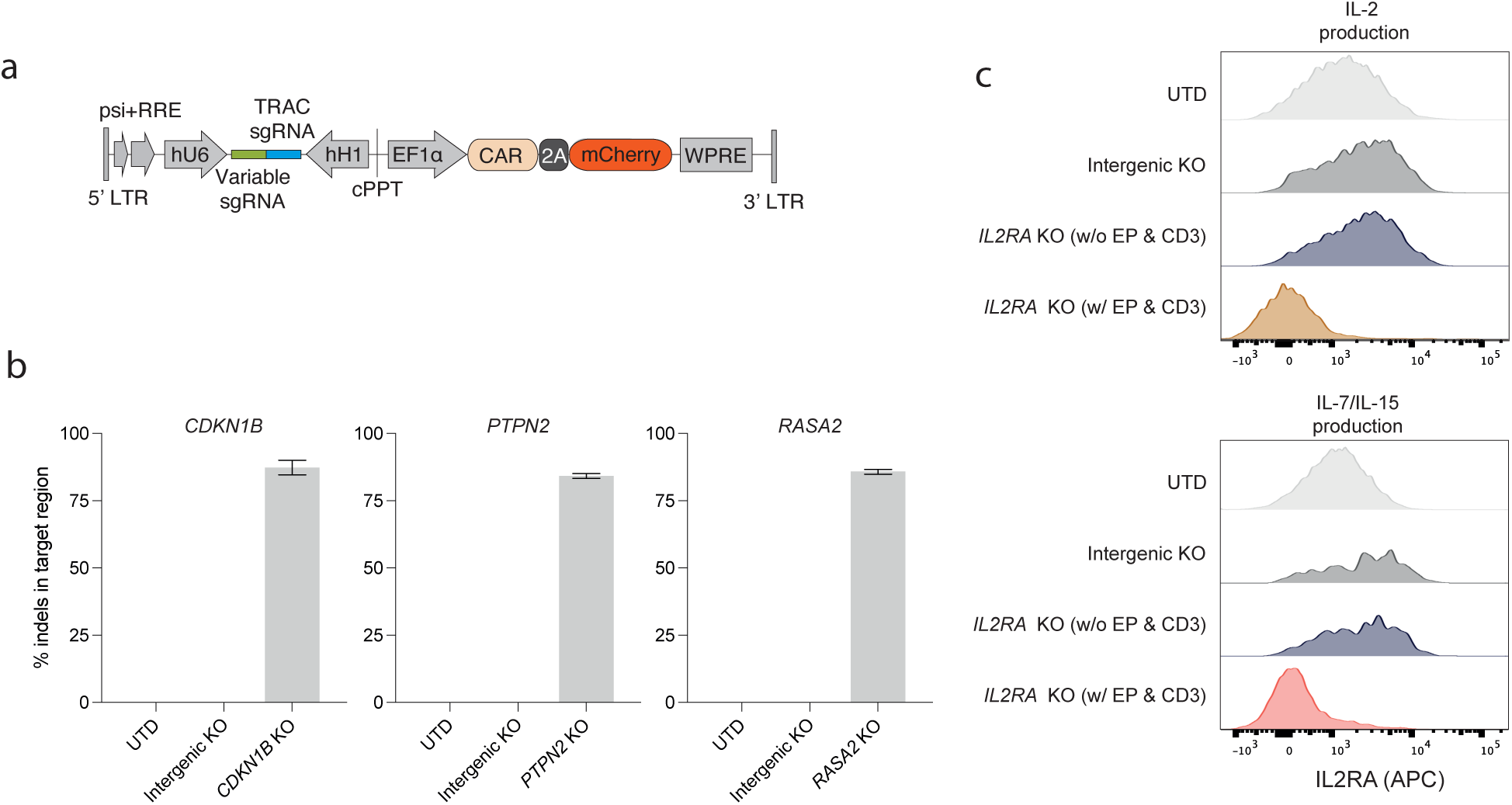
Generation and validation of knockout CAR T cells. **a.** Construct design for the 4-1BB BCMA CAR (pCAR) and double-guide cassette (pGuide) containing individual gene sgRNAs (variable sgRNA): *CDKN1B*, *IL2RA*, *PTPN2*, and *RASA2*. **b.** Quantification of insertions/deletions (indels) in sgRNA target region by next generation sequencing. Data represents target sgRNA 1 and 2 for *CDKN1B*, *PTPN2*, and *RASA2*, respectively; presented as mean +/-SEM. **c.** *IL2RA* expression by flow cytometry, with *IL2RA* KO CAR T cells (before electroporation/CD3 negative selection and after) compared to UTD and intergenic control KO (both IL-2 and IL-7/IL-15-produced, respectively). UTD = untransduced T cells.

**Extended Data Figure 4.**
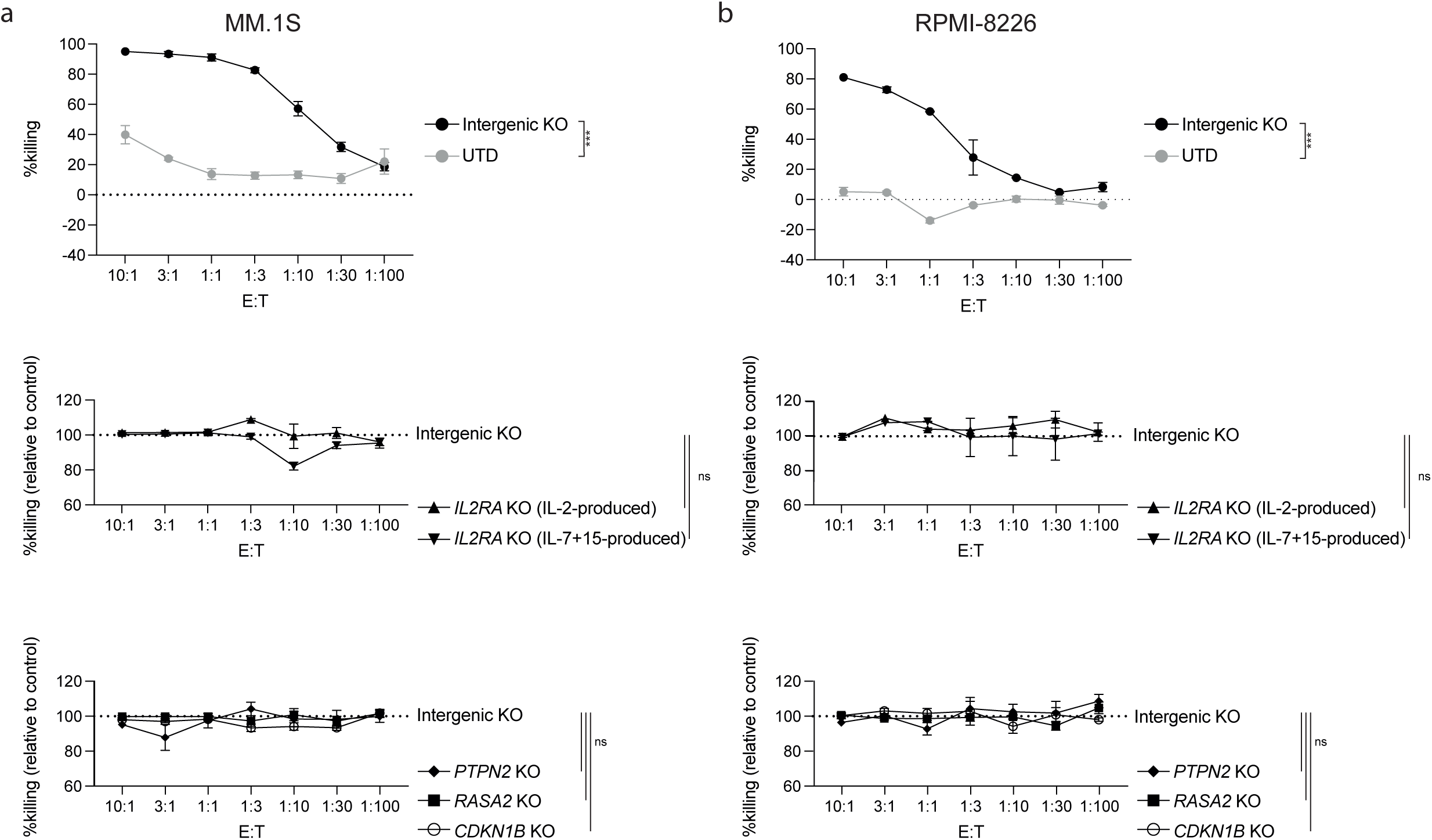
In vitro cytotoxicity of knockout CAR T cells. **a.** 16-hour luciferase-based killing assay of knockout CAR T cells co-cultured with MM.1S tumor cells, in different effector to target (E:T) ratios (from 10:1 to 1:100). Upper panels display comparison of intergenic KO CAR T cells to UTD. Middle and lower panels show relative killing of *IL2RA* KO (IL-2-or IL-7/IL-15-produced; middle panel) or *CDKN1B* KO, PTPN2 KO, and RASA2 KO (lower panel) compared to intergenic KO CAR T cells. Data represent technical replicates from CAR T cells generated from two normal donors (ND116, ND202). Data presented as mean +/-SEM. Statistical significance was measured by two-way ANOVA with Tukey’s multiple comparison test. UTD = untransduced T cells. ***p < 0.001, and ns = non significant.

**Extended Data Figure 5.**
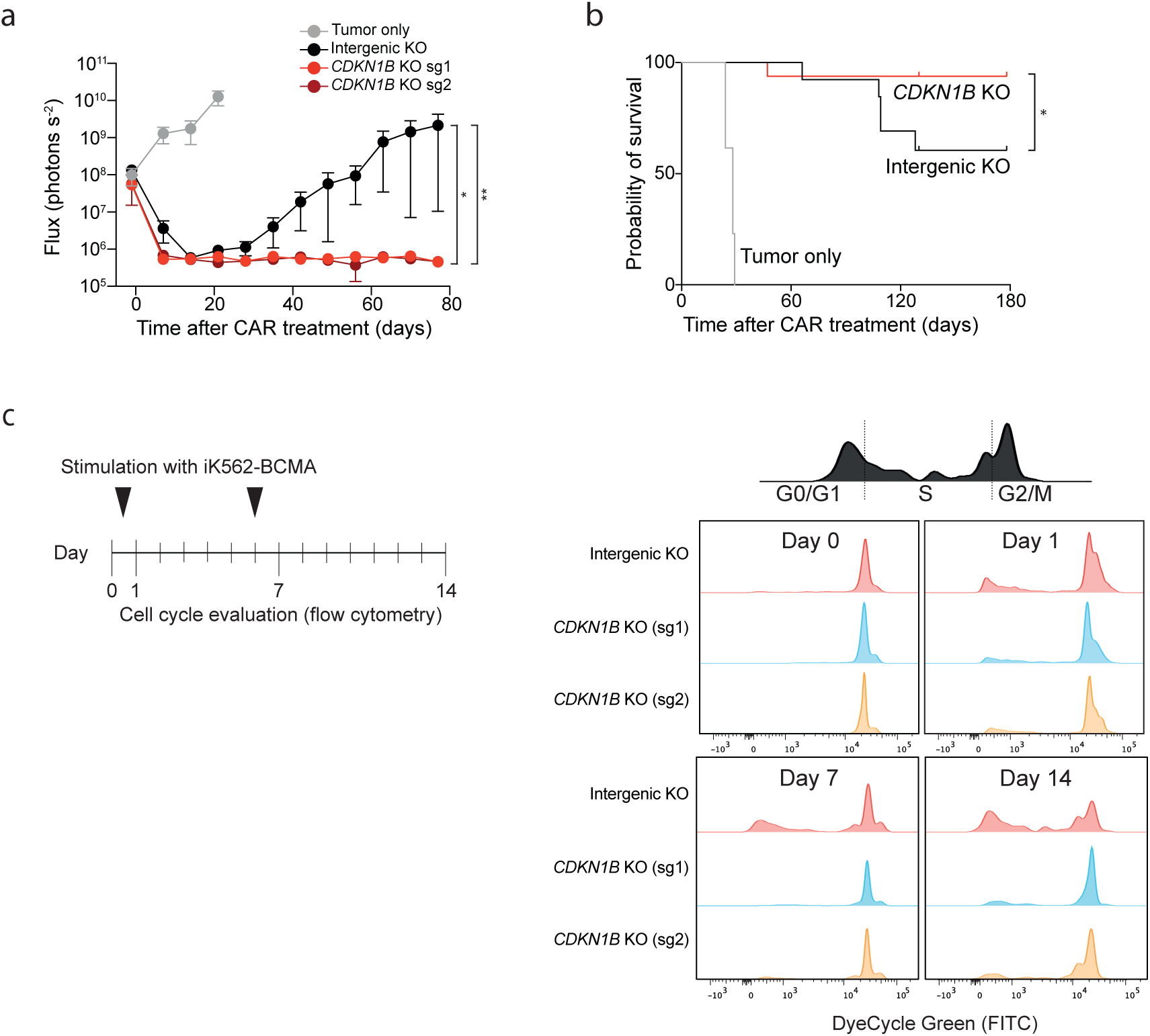
*CDKN1B* ablation increases the efficacy of BCMA CAR T cells in vivo. **a.** MM.1S tumor burden as measured by bioluminescent imaging (BLI) of mice treated with intergenic control KO, or *CDKN1B* KO BCMA CAR T cells. NSG mice were injected intravenously with 1E6 MM.1S followed by transfer of CAR T cells 21 days later. n=3 mice per group from one healthy donor (ND202). Data presented as mean +/-SEM. Statistical significance was measured compared to the intergenic KO CAR T cell treated group at day 77 (*CDKN1B* KO, *PTPN2* KO) as measured by two-way ANOVA with Tukey’s multiple comparison test. **b.** Overall survival of mice treated with intergenic control KO or *CDKN1B* KO BCMA CAR T cells. n=13 for tumor only and intergenic control KO CAR T cell treated animals and n=16 for *CDKN1B* KO CAR T cell treated animals. Data were combined from multiple experiments using T cells from two healthy human donors (ND202, ND116). Statistical significance was measured by log-rank (Mantel–Cox test) for Kaplan–Meier curves. **c.** Comparison of cell cycle (G0/G1, S, G2/M) between *CDKN1B* KO and intergenic control KO CAR T cells. Stimulation with irradiated K562-BCMA tumor cells was conducted on day 0 and day 6, with flow cytometry measurements on day 0 (before stimulation), day 1, day 7, and day 14 (schematic overview, left). *p < 0.05, and **p < 0.01.

## Extended Data Tables

**Extended Data Table 1. Mario sgRNA library list of genes and sgRNA sequences**

**Extended Data Table 2. Results of BCMA Mario-CAR T cell screen using IL-2 during in vitro production**

**Extended Data Table 3. Results of BCMA Mario-CAR T cell screen using IL-7/IL-15 during in vitro production**

**Extended Data Table 4. Perturb-seq sgRNA library list of genes and sgRNA sequences**

**Extended Data Table 5. Differential genes driving scRNAseq clusters**

**Extended Data Table 6. Pseudo-bulk Hallmark GSEA analysis results from perturb-seq experiment**

**Extended Data Table 7. Differential gene expression analysis comparing *CDKN1B* KO and intergenic KO CAR T cells from bulk RNA sequencing**

**Extended Data Table 8. Hallmark GSEA analysis results comparing *CDKN1B* KO and intergenic KO CAR T cells from bulk RNA sequencing**

**Extended Data Table 9. GSEA analysis of T cell phenotype gene sets comparing *CDKN1B* KO and intergenic KO CAR T cells from bulk RNA sequencing**

